## Supplementary figure 1 for "A divergent cyclin/cyclin-dependent kinase complex controls the atypical replication of a malaria parasite during gametogony and transmission"

### Figure 1 - supplement 1

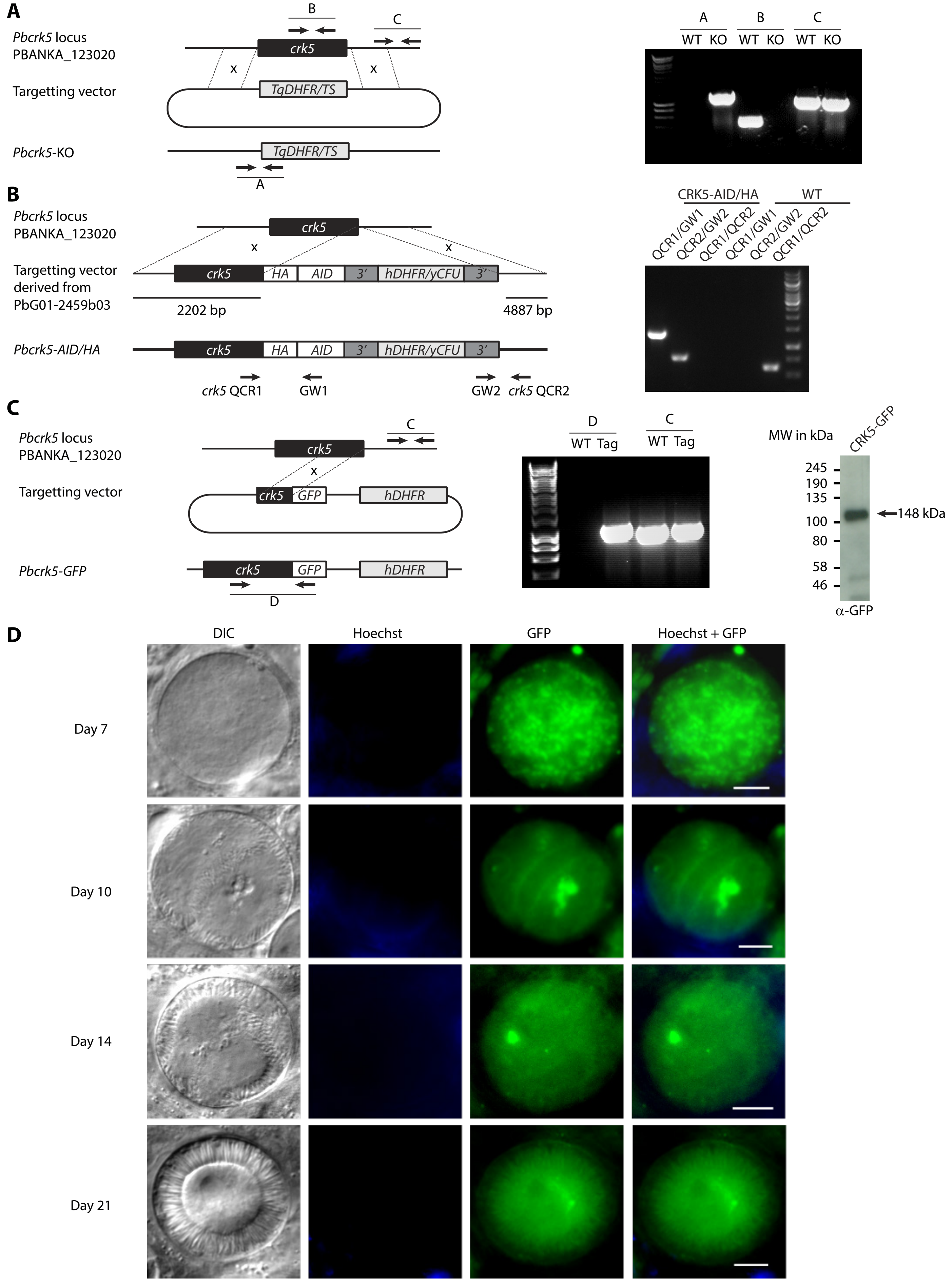


**Figure S1. Generation of CRK5-KO, CRK5-AID/HA, and CDKrs-GFP transgenic lines and CRK5-GFP expression in oocysts.**
A-C. Genetic modification vectors and genotyping data for CRK5-KO (A), CRK5-AID/HA (B), and CRK5-GFP (C). Oligonucleotides used for PCR genotyping are indicated and agarose gels for corresponding PCR products from genotyping reactions are shown. C. Western blot detection of CRK5-GFP.

D. CRK5-GFP expression and localisation in midgut oocyst at day 7, 14, 17, and 21 post-infection. Scale bar = 10 µm.
