## Supplementary figure 2 for "A divergent cyclin/cyclin-dependent kinase complex controls the atypical replication of a malaria parasite during gametogony and transmission"

### Figure 2 - supplement 1

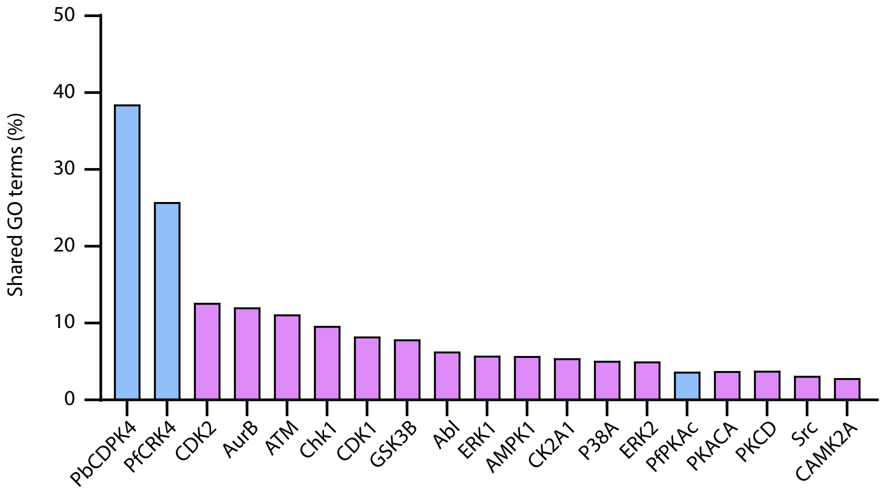


**Figure S2. Percentage of GO terms shared between CRK5 and a set of *Plasmodium* and human kinases.**
For CRK5 GO terms, proteins with a >2 fold change (*p*-value <0.05) in the CRK5-KO were retained.
