## Supplementary figure 3 for "A divergent cyclin/cyclin-dependent kinase complex controls the atypical replication of a malaria parasite during gametogony and transmission"

### Figure 3 - supplement 1

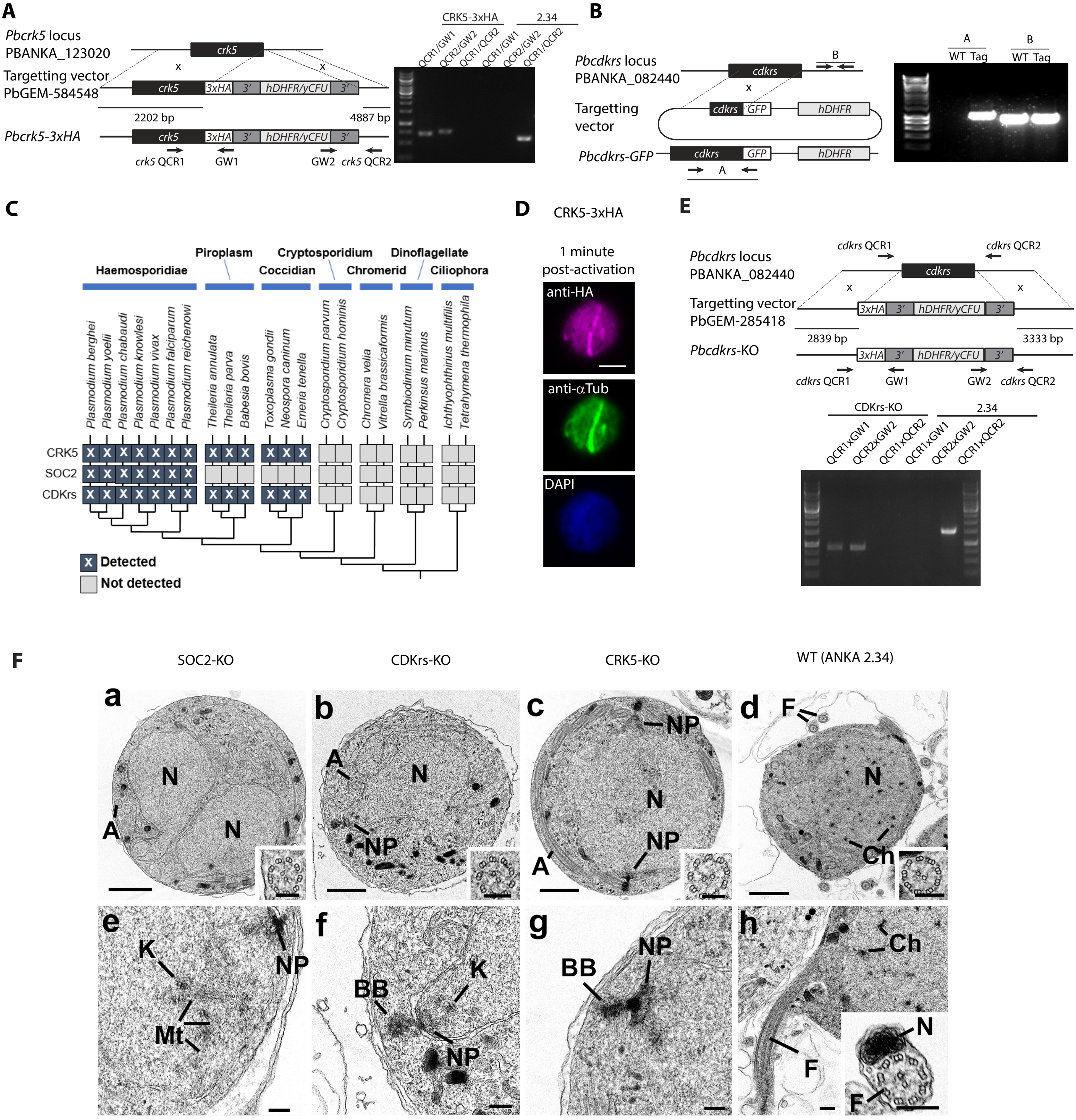


**Figure S3. Identification and characterisation of the CSC complex components.**

A-B. Genetic modification vectors and genotyping data for CRK5-HA (A) and CDKrs-GFP (B). Oligonucleotides used for PCR genotyping are indicated and agarose gels for corresponding PCR products from genotyping reactions are shown.

C. Phylogenetic distribution of the CSC complex components and of Plasmodium CRKs in selected alveolate genomes (see Methods). Branch support values evaluated with aBayes are shown on branches.

D. Localisation of CRK5-HA in 1 min activated gametocytes. Scale bar = 5 µm.

E. Genetic modification vectors and genotyping data for CDKrs-KO.

F. Electron micrographs of the male gametocytes of the SOC2-KO (a, e), CDKrs-KO (b, f), CRK5-KO (c, g) and wild type (d,h) parasites at 15 min pa. The three KO parasites consist of predominately early stages (95-100%) while the WT showed a number of late stages (~40%) with exflagellation and free microgametes. Bars represent 1 µm in a-d and 100 nm in all other micrographs. (a to c) SOC2-KO, CDKrs, and CRK5 male gametocytes showing the early stage of development with a lobed nucleus (N) and cytoplasmic axonemes (A). Inserts. Detail of cross-sectioned axonemes. (d) WT showing a late stage in development with nucleus (N) containing small clumps of condensed chromatin (Ch). Note the free microgamete flagella (F). Insert. Cross-sectioned axoneme. (e) Detail of a similar stage parasite to that in (a) showing a nuclear pole (NP) but note the disorganized arrangement of spindle microtubules (Mt) and attached kinetochores (K). (f) Detail of the nucleus in (b) showing the nuclear pole (NP) with a closely associated basal body (BB) in the cytoplasm. K – Kinetochore. (g) Detail of a similar stage to that in (c) showing the nuclear pole (NP) and associated basal body (BB). (h) Detail of a similar parasite to that in (d) undergoing exflagellation showing the flagellum (F) protruding from the microgametocyte. Ch – condensed chromatin. Insert. Cross section through a free microgamete showing the flagellum (F) and electron dense nucleus (N).
