## Supplementary figure 4 for "A divergent cyclin/cyclin-dependent kinase complex controls the atypical replication of a malaria parasite during gametogony and transmission"

### Figure 4 - supplement 1

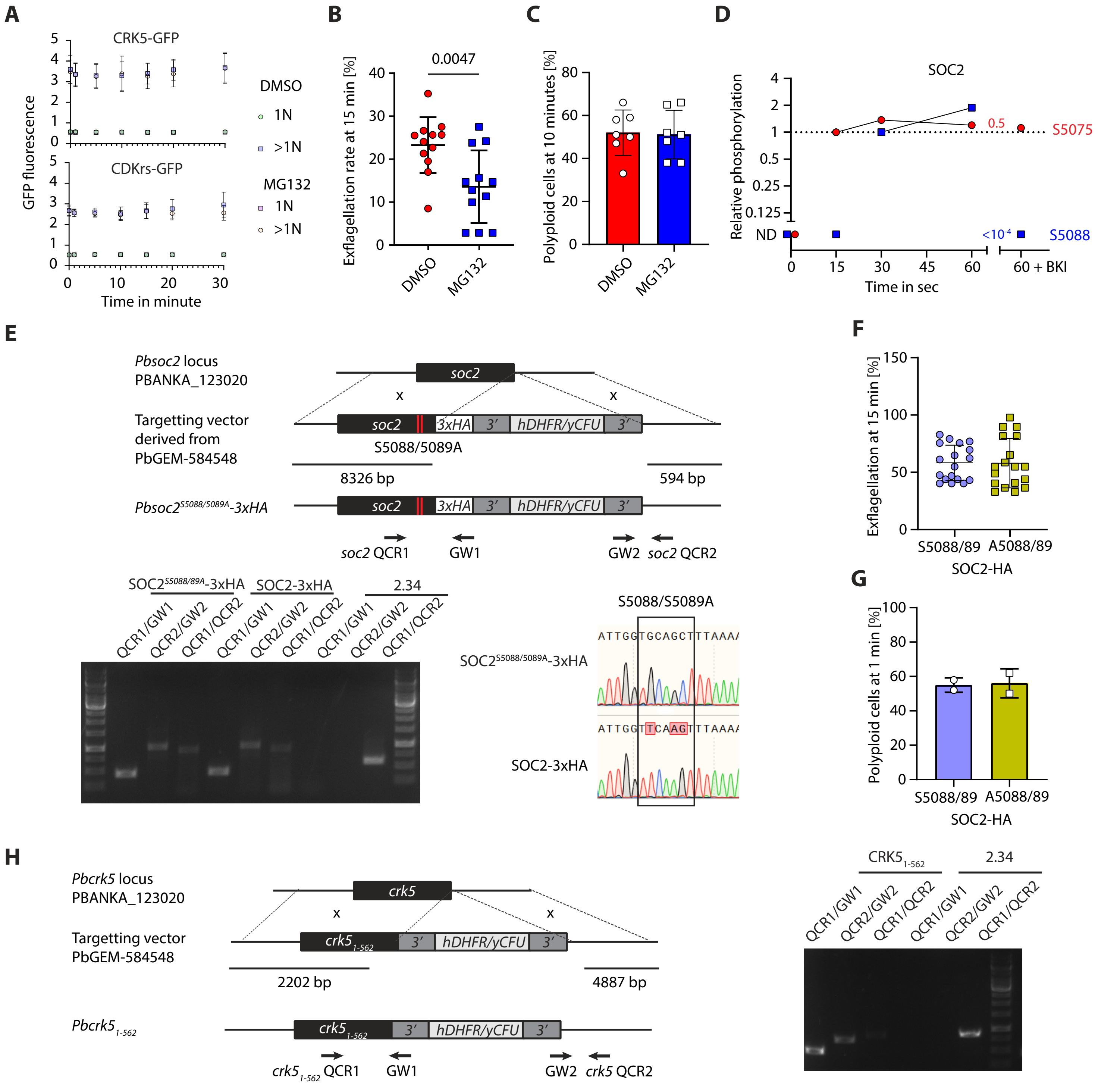


**Figure S4. Study of the CSC complex regulation.**

A. Flow cytometry analysis of CRK5-GFP and CDKrs-GFP fluorescence over the course of gametogony in haploid or multiploid (>1N) cells in the presence or absence of 1 µM of MG132 a proteasome inhibitor (error bars show standard deviation from the mean; 3 independent infections; two-way ANOVA).

B. Pre-treatment of gametocytes with 1 µM MG132 leads to reduced exflagellation (error bars show standard deviation from the mean; technical replicates from 3 independent infections; unpaired two-tailed t-test).

C. Treatment with 1 µM MG132 has no effect on the number of male gametocytes replicating their DNA. The proportion of male gametocytes undergoing DNA replication was determined at 1 min pa and is expressed as a percentage of cells that are polyploid (>1 nucleus) (error bars show standard deviation from the mean; technical replicates from 3 independent infections; unpaired two-tailed t-test).

D. SOC2 phosphorylation levels show that S5088 is phosphorylated in a CDPK4-dependent manner 15 sec after activation. ND = not detected. BKI = CDPK4 inhibitor

E. Genetic modification vector and genotyping data for SOC2$^{S5088/5089A}$ mutagenesis. Oligonucleotides used for PCR genotyping are indicated and agarose gels for corresponding PCR products from genotyping reactions are shown.

F. S5088/5089A substitution in SOC2-3xHA does not impair exflagellation (error bars show standard deviation from the mean; technical replicates from 2 independent infections; two-way ANOVA).

G. There is no reduction in the number of male gametocytes replicating their DNA in SOC2$^{S5088/5089A}$-3xHA gametocytes. The proportion of male gametocytes undergoing DNA replication was determined at 1 min pa and is expressed as a percentage of cells that are polyploid (>1N) (error bars show standard deviation from the mean; from 2 independent infections; two-way ANOVA).

H. Genetic modification vector and genotyping data for CRK5 mutagenesis. Oligonucleotides used for PCR genotyping are indicated and agarose gels for corresponding PCR products from genotyping reactions are shown.
