## Supplementary figures and images for "A divergent cyclin/cyclin-dependent kinase complex controls the atypical replication of a malaria parasite during gametogony and transmission"

### Figure 1 source data 2

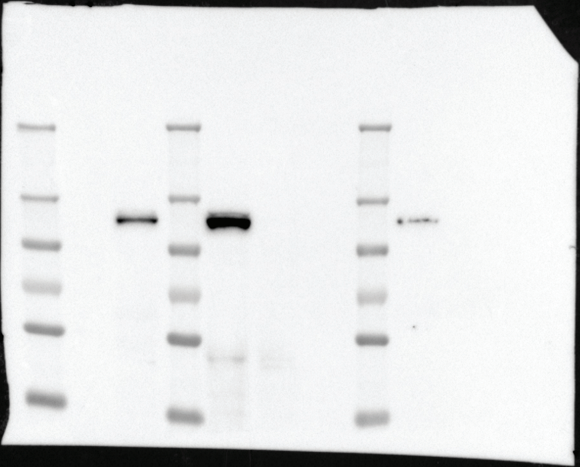

### Figure 1 source data 3

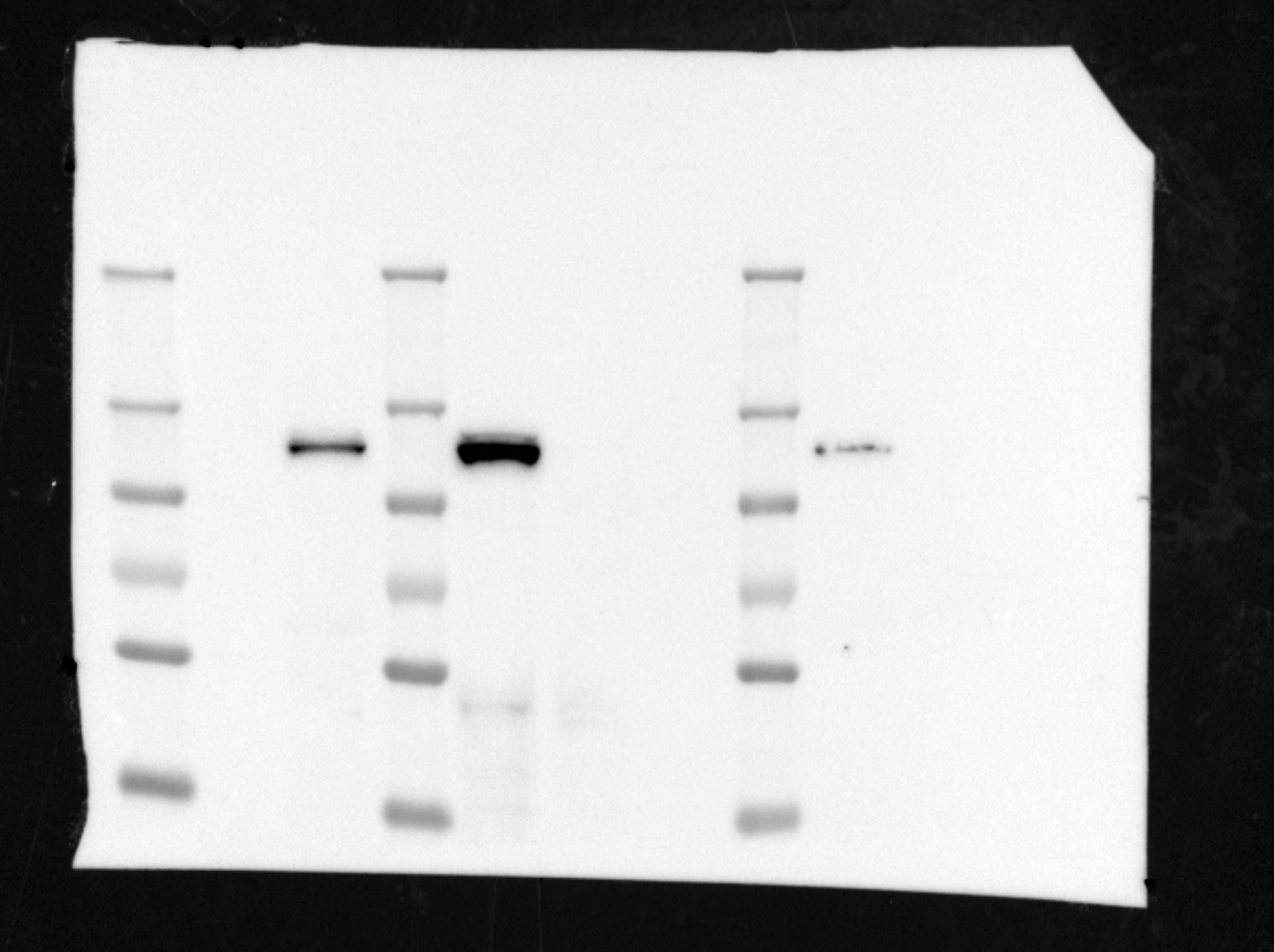

### Figure 1 source data 3

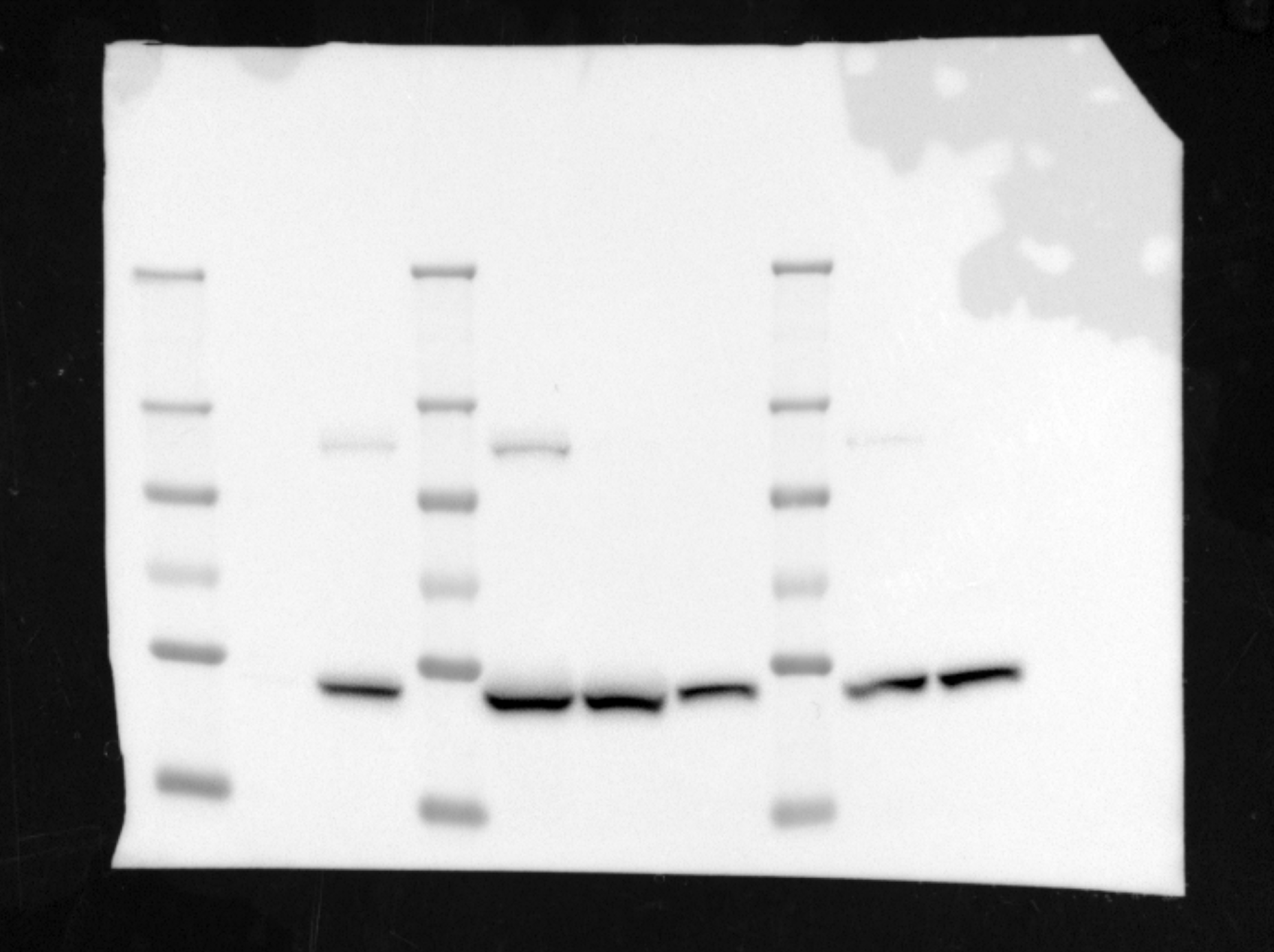

### Figure 1 source data 4

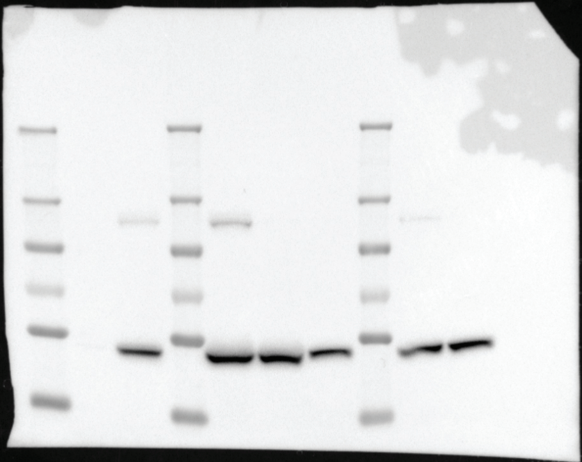

### Figure 4 source data 1

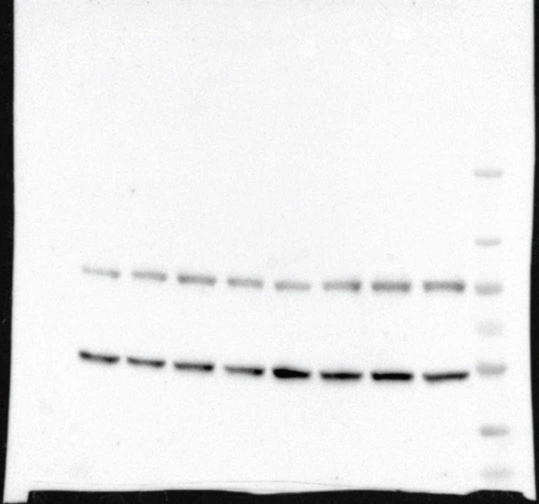

### Figure 4 source data 2

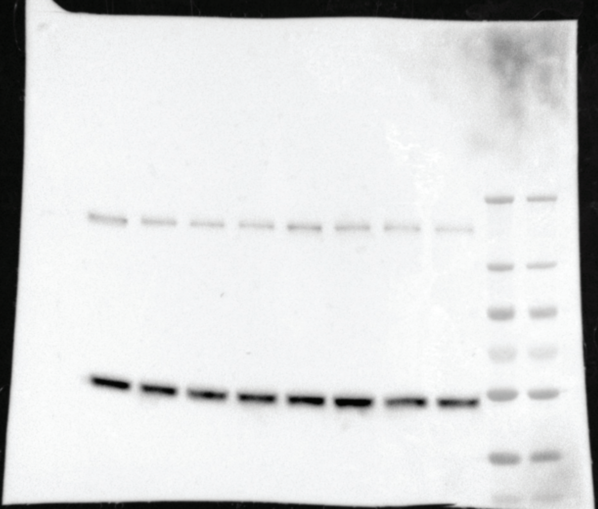

### Figure 4 source data 2

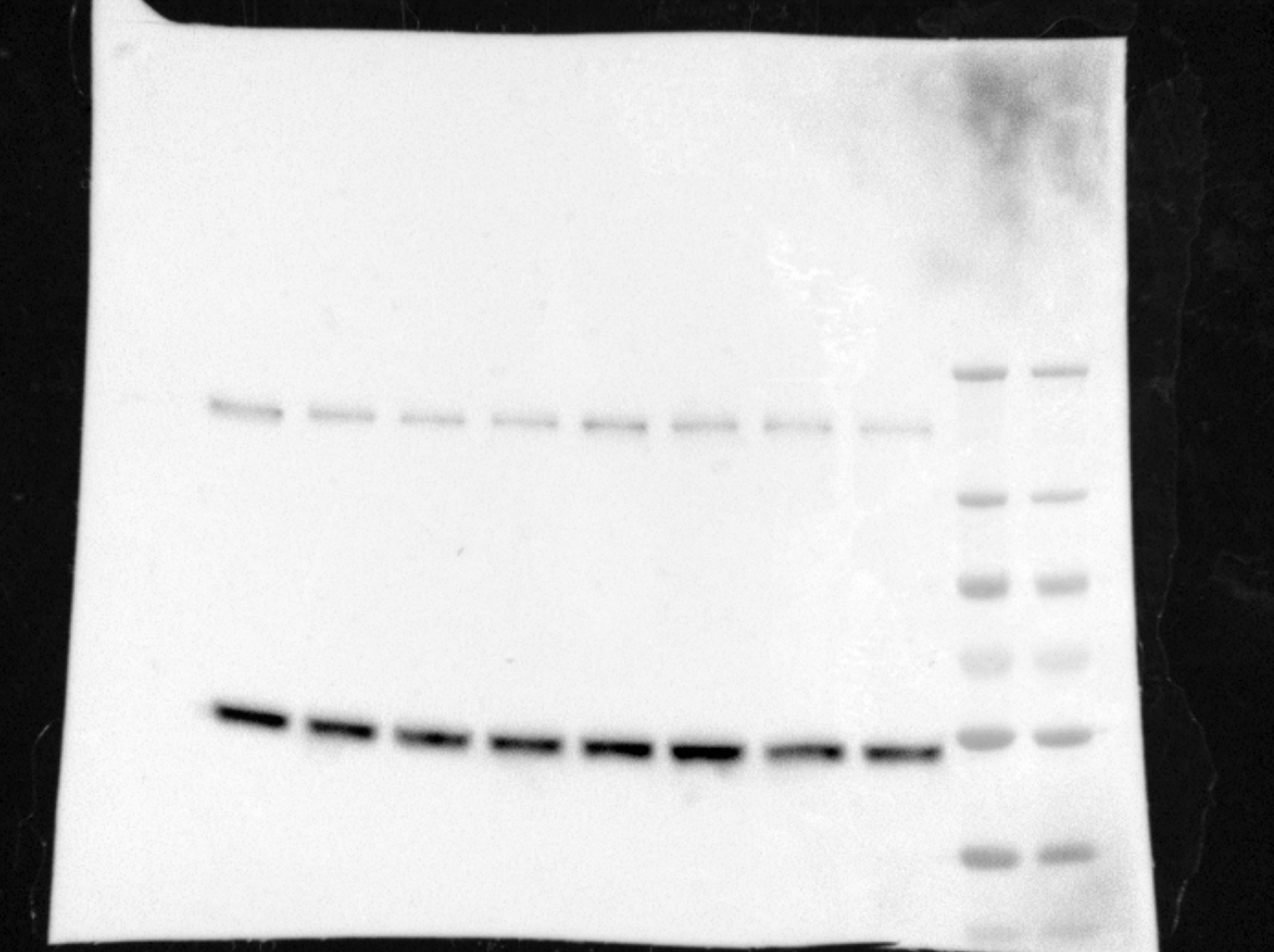

### Figure source data 1

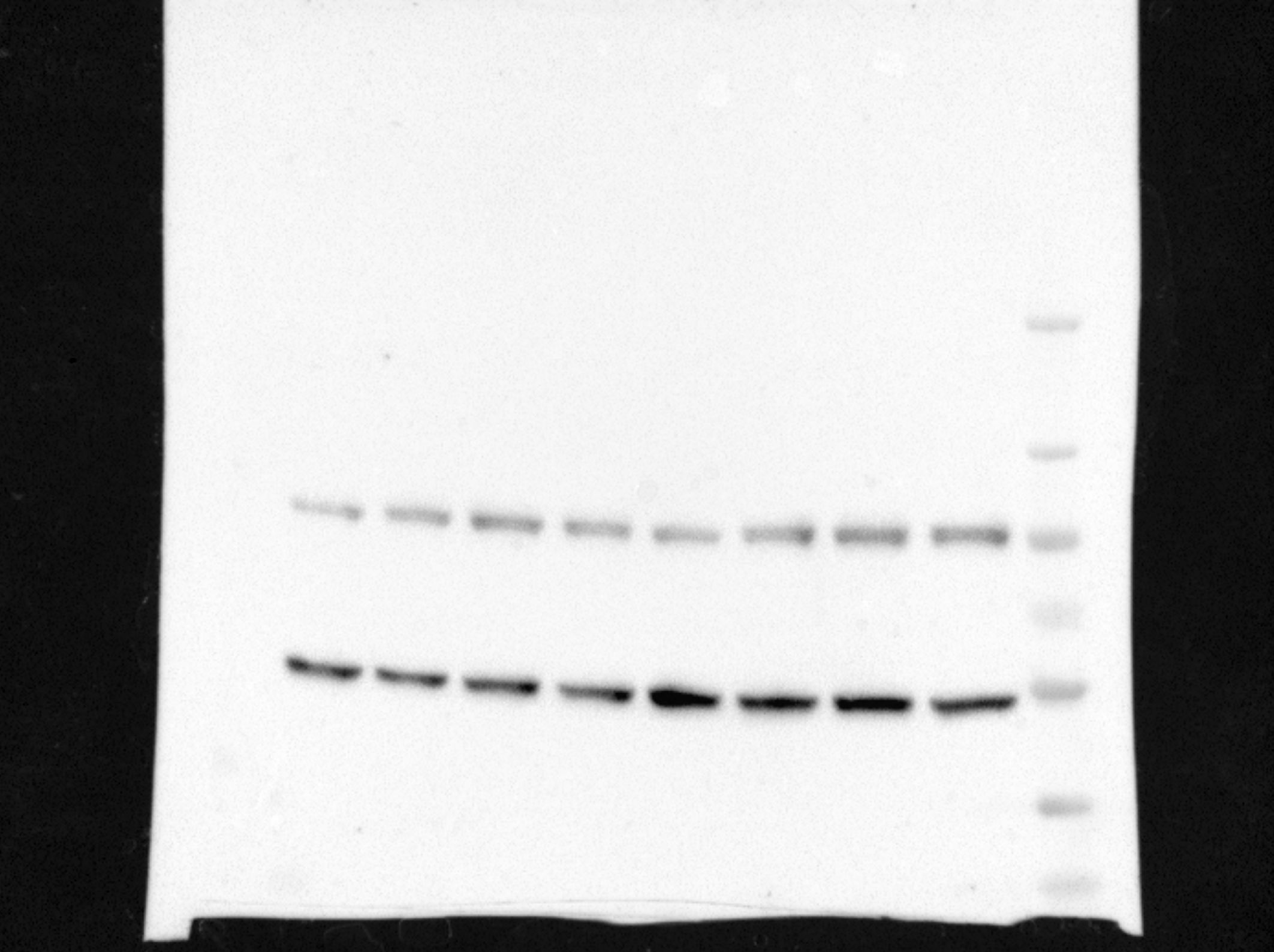
